# Subjective value, not a gridlike code, describes neural activity in ventromedial prefrontal cortex during value-based decision-making

**DOI:** 10.1101/759951

**Authors:** Sangil Lee, Linda Q. Yu, Caryn Lerman, Joseph W. Kable

## Abstract

Across many studies, ventromedial prefrontal cortex (vmPFC) activity has been found to correlate with subjective value during value-based decision-making. Recently, however, vmPFC has also been shown to reflect a hexagonal gridlike code during navigation through physical and conceptual space. This raises the possibility that the subjective value correlates previously observed in vmPFC may have actually been a misconstrued gridlike signal. Here, we first show that, in theory, a hexagonal gridlike code of two-dimensional attribute space could mimic vmPFC activity previously attributed to subjective value. However, using fMRI data from a large number of subjects performing an intertemporal choice task, we show clear and unambiguous evidence that subjective value is a better description of vmPFC activity than a hexagonal gridlike code. In fact, we find no significant evidence at all for a hexagonal gridlike code in vmPFC activity during intertemporal choice. This result limits the generality of gridlike modulation as description of vmPFC activity. We suggest that vmPFC may flexibly switch representational schemes so as to encode the most relevant information for the current task.

---

Many studies in decision neuroscience have identified a critical role for the ventromedial prefrontal cortex (vmPFC) in decision-making. Specifically, neural activity in the vmPFC correlates with the subjective value of expected or experienced outcomes across a wide variety of decision-making tasks (Bartra, McGuire, & Kable, 2013; Clithero & Rangel, 2013; Levy & Glimcher, 2012). Neural correlates of subjective value have been found in vmPFC using both fMRI in humans as well as single cell recording in non-human animals (Howard, Gottfried, Tobler, & Kahnt, 2015; Kable & Glimcher, 2007; McNamee, Rangel, & O’Doherty, 2013; Strait, Blanchard, & Hayden, 2014; Yamada, Louie, Tymula, & Glimcher, 2018). One straightforward interpretation of these results is that vmPFC encodes the subjective value of potential outcomes, which is then used to make choices between outcomes (Kable & Glimcher, 2009; Rich & Wallis, 2016).

However, recent human neuroimaging studies have found that a similar area of vmPFC also serves a different function: encoding representational maps that enable navigation through physical and conceptual spaces (Constantinescu, O’Reilly, & Behrens, 2016; Doeller, Barry, & Burgess, 2010; Jacobs et al., 2013). Both intracortical recordings and fMRI studies of humans navigating through virtual arenas have shown that activity in vmPFC is modulated in a hexagonal manner by the direction of travel, which is a pattern characteristic of grid cells (Bao et al., 2019; Doeller et al., 2010; Jacobs et al., 2013; Nau, Navarro Schröder, Bellmund, & Doeller, 2018). Grid cells were first discovered in entorhinal cortex (ERC) during spatial navigation and provide an efficient representation of two-dimensional space (Behrens et al., 2018; Doeller et al., 2010; Hafting, Fyhn, Molden, Moser, & Moser, 2005). More recently, fMRI signatures of this hexagonal gridlike code have been observed in vmPFC during navigation in a purely conceptual space (Constantinescu et al., 2016). Specifically, stimuli in that study were defined along two dimensions, and when subjects imagined a stimulus transforming through the conceptual space defined by those two dimensions, activity in vmPFC showed a similar response pattern as that observed during two-dimensional spatial navigation.

These recent results have led some to speculate that these hexagonal gridlike codes might serve as a general account for the role of vmPFC in complex cognition, including decision-making (Bellmund, Gärdenfors, Moser, & Doeller, 2018). This is conceivable as many choices in decision-making are arguably between options that differ along (at least) two dimensions – for example, gambles that differ in risk and payoff, foods that differ in health and taste, or goods that differ in quality and price. Might vmPFC activity during decision-making reflect navigation through a conceptual space defined by these attribute dimensions? If true, hexagonal gridlike codes could provide a general account of vmPFC function across multiple domains.

Here we put this idea to the test: does the vmPFC exhibit hexagonal gridlike modulation during decision making, thereby reflecting conceptual navigation through attribute space? We first establish the theoretical plausibility of this idea by showing that a hexagonal gridlike modulation signal can be highly correlated with subjective value and thus could be mistaken for it. We then empirically test if BOLD activity in vmPFC during value-based decision-making can be explained by a hexagonal gridlike code, using a large existing intertemporal choice dataset (Kable, Caulfield, Falcone, McConnell, Bernardo, Parthasarathi, Cooper, Ashare, Audrain-McGovern, Hornik, et al., 2017). Across three different analyses, we show unambiguously that vmPFC activity in the intertemporal choice task is better described by a subjective value signal than by a hexagonal gridlike modulation. This finding limits the generality of hexagonal grid representations in vmPFC. Instead, vmPFC may flexibly switch representational schemes so as to encode the most pertinent information to the task at hand.

## Methods

### Dataset

We used the fMRI dataset from Kable et al. (2017), as its large number of subjects permits a high-powered test between a subjective value and hexagonal grid code during decisions that involve a tradeoff between two choice attributes. The dataset is online at OpenNeuro.org and will be made public upon acceptance of the manuscript. Participants completed two sessions in which they performed an intertemporal choice task in the scanner 10 weeks apart. In each scan session, participants made 120 binary choices between a smaller immediate reward and a larger later reward. The smaller immediate reward was held constant at $20 today while the larger later reward varied in amount (*A*: $21 ∼ $85) and delay (*D*: 20 days ∼ 180 days) from trial to trial. On each trial, the amount and delay of the larger later option was displayed on the screen, while the constant immediate option was not displayed. Participants used a button pad to indicate whether they would accept the larger delayed option shown on the screen or to reject it in favor of the smaller immediate option, which was not shown on the screen. This design simplifies the identification of neural correlates of subjective value, as only the subjective value of the delayed option needs to be considered; since the immediate option is fixed, the sum, difference or ratio of the subjective values of the two options are all linearly related to the subjective value of the delayed option.

The two hypotheses provide different predictions of the expected signal in this task (**Fig. 1**). Subjective values depend on how individual subjects weight the two attribute dimensions, but all subjects prefer larger magnitudes and smaller delays, leading to the highest response in one corner of the two-dimensional attribute space. For hexagonal grid modulation, the relevant conceptual navigation in this task involves “traversing” between the two options being considered. A gridlike code would lead to peaks in activity when navigating along hexagonally symmetric directions spaced sixty degrees apart, with the angle of these peak directions (the grid angle) being the one free parameter. For example, a person with a grid-angle at 0 degrees would have activity peaks at 0 and 60 degrees of traversing angle and troughs at 30 and 90 degrees, while another person with a grid angle at 30 degrees would have peaks at 30 and 90 degrees and troughs at 0 and 60 degrees.

**Figure 1.**
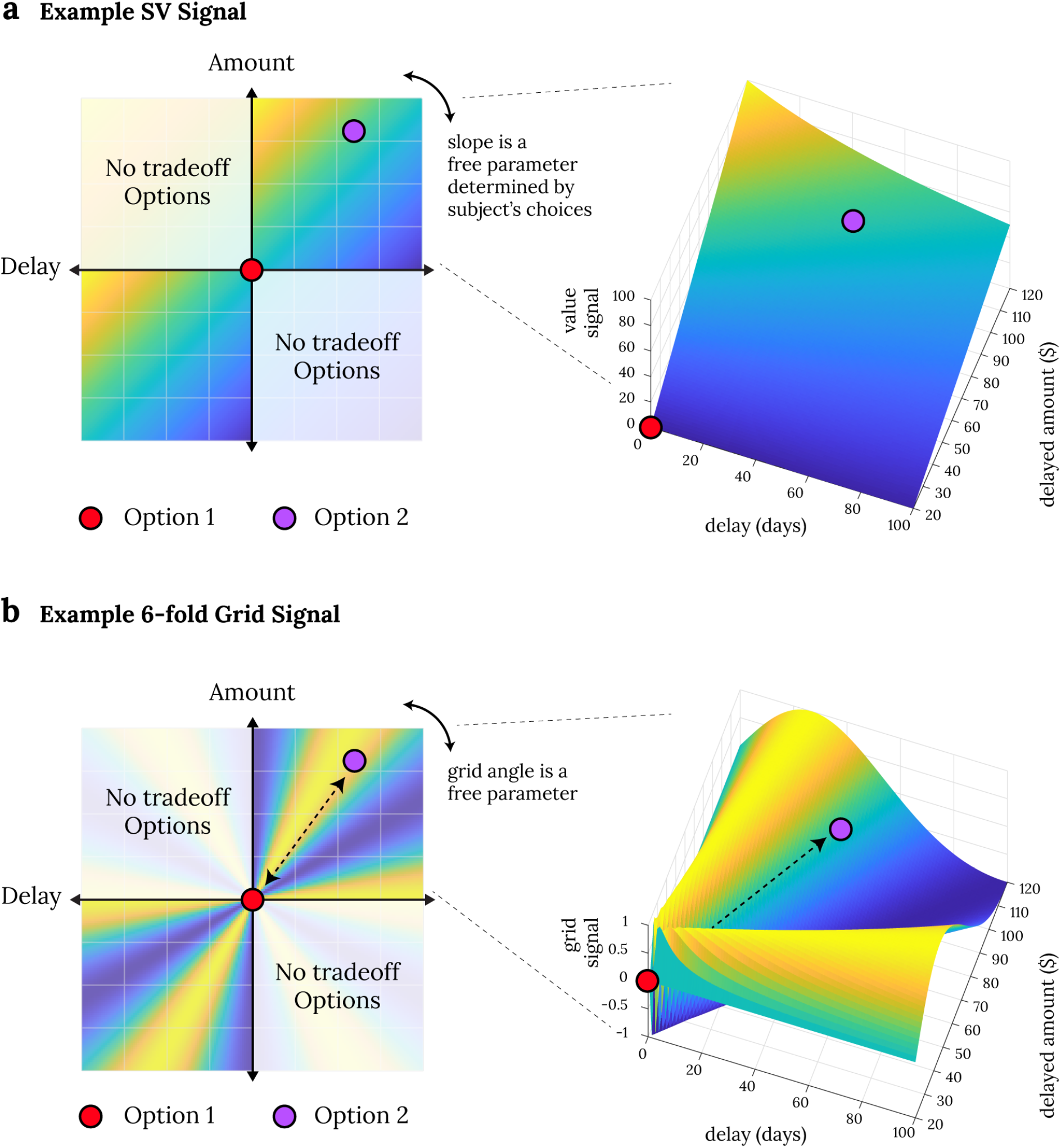
Example SV signal and hexagonal gridlike signal in two attribute space. Panel **a** shows an example subjective value signal when comparing two options that vary in amount and delay. In the choice task used here, an SV signal will vary depending on the subjective value of the variable delayed option. Panel b shows an example hexagonal gridlike signal when mentally traversing between two points (shown in example is a grid angle of 0 degrees, such that the peak activity occurs when traversing at 0 and 60 degrees; this grid angle may vary from person to person). In the choice task used here, a hexagonal gridlike signal will vary depending on the traversing angle between the two options.

Of the 160 participants that completed session 1, we excluded participants with any missing runs (n = 6), too much head movement (any run out of 4 runs with >5% of mean image displacements greater than 0.5mm; n = 3), more than 3 missing trials per run for two or more runs (n = 2), entirely one-sided choice such that the participant always chose the immediate or the delayed option (n = 3), and one participant who expressed knowledge of his experimental condition in the original study. This resulted in the final sample size of 145 participants for session 1. Of these participants, only 114 completed session 2, from which we also excluded those with missing runs (n = 3), too much head movement (n = 2), too many missing trials (n = 2), or entirely one-sided choices (n = 5). This gave us a total of 102 participants for session 2. In total, 145 participants’ data were used for session 1 and 102 participants’ data were used when analyzing across both sessions.

Participants were scanned with a Siemens 3T Trio scanner with a 32-channel head coil. T1-weighted anatomical images were acquired using an MPRAGE sequence (T1 = 1100ms; 160 axial slices, 0.9375 x 0.9375 x 1.000 mm; 192 x 256 matrix). T2*-weighted functional images were acquired using an EPI sequence with 3mm isotropic voxels, (64 x 64 matrix, TR = 3,000ms, TE = 25ms; tilt angle = 30°) involving 53 axial slices with 104 volumes. B0 fieldmap images were also collected for distortion correction (TR = 1270ms, TE = 5 and 7.46ms). The datasets were preprocessed via FSL FEAT (FMRIB fMRI Expert Analysis Tool). Functional images were skull stripped with BET (FMRIB Brain Extraction Tool), motion corrected and aligned with MCFLIRT (FMRIB Linear Image Restoration Tool with Motion Correction), spatially smoothed with a FWHM 9mm Gaussian Kernel, and high pass filtered (104sec cutoff). Registration was performed with FNIRT with warp resolution of 20mm (FMRIB’s Non-linear Image Registration Tool) to a 2mm MNI standard brain template.

### Could a hexagonal grid signal be mistaken for a subjective value signal?

Before any empirical analysis, we sought to show that, in theory, hexagonal gridlike modulation could mimic or account for activity correlated with subjective value during decision making. To do this, we simulated a subjective value signal at a given discount rate for a range of amounts and delays, and then estimated the best-fitting gridlike modulation for this signal. For a given discount rate, the subjective values of delayed monetary outcomes were calculated using the hyperbolic model:

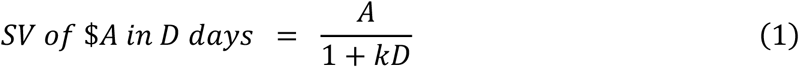

where *k* is the individual discount rate. The amount *A* varied from 20 to 80 in increments of 2 (31 levels) and the delay *D* varied from 20 to 180 in increments of 5 (33 levels) resulting in a total of 1023 subjective values for a given *k*. This 1023-element vector (*SV*) was then regressed against two hexagonal grid modulation regressors:

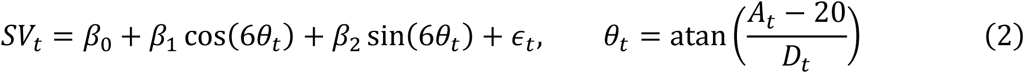

The model contains a linear combination of the sine and cosine of the trajectory angle *θ* with 60° periodicity. The trajectory angle *θ* is taken as the angle between the abscissa and the traversing line between the immediate option ($20 now) and the delayed option ($*A* in *D* days) in the two-dimensional space defined by the amount and delay attributes (e.g., **Fig. 1d**). The linear sum of cosine and sine function acts as a phase shift (i.e., generally: *p*cosθ + *q*sinθ = *k*cos(θ + ϕ)) such that we can identify the best phase (i.e., grid angle) of the cosine modulation without using a non-linear regression. After fitting the regression, we calculated the Pearson correlation between *SV* and the fitted signal to assess the similarity between the two. This procedure was repeated for 51 levels of *k* whose base-10 log ranged from −5 (i.e., *k* = 0.00001: very patient) to 0 (i.e., *k* = 1: very impatient) in 0.1 increments.

### fMRI analysis – voxelwise GLM

In empirical fMRI data, we first performed two GLMs in session 1 data to test for activity that was correlated with subjective value or hexagonal grid signals. For the subjective value GLM, we used two regressors: an event regressor that modeled average activity of all trials, and second regressor that modeled activity modulated by subjective value. The subjective value of the delayed reward was estimated by fitting a hyperbolic discounting model to choice data using a logit choice model (*A* is the delayed amount, *D* is the delay, *k* is the discount rate, and *β* is the scaling factor):

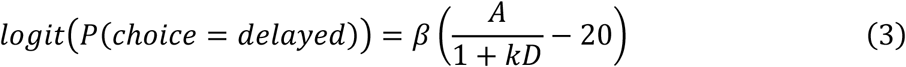

Regions correlated with subjective value were identified by performing a sign-flipping permutation test on individual beta images of the subjective value regressor. For the gridlike modulation GLM, we used three regressors: an event regressor and two hexagonal grid angle regressors (*n* = 6):

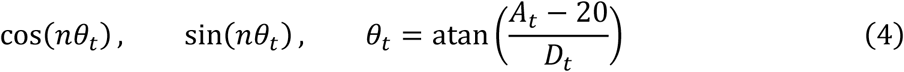

Regions correlated with hexagonal grid signal were identified by first transforming the *F*-statistic of the two hexagonal grid regressors to a *z*-statistic and then by performing a sign-flipping permutation test on the *z*-transformed *F*-stat images (Constantinescu et al., 2016).

### fMRI analysis – model comparison

We tested whether vmPFC activity is better described by a subjective value or hexagonal gridlike signal in three different ways. First, and most straightforward, we performed a model comparison between the subjective value GLM and the hexagonal modulation GLM. We compared the Akaike Information Criterion (AIC) scores of the two GLM models in each of four ROIs: vmPFC and ventral striatum ROIs from the Bartra et al. (2013) meta-analysis of subjective value, and two spherical ROIs (radius = 2 voxels) from the peak activation coordinates reported by Constantinescu et al. (2016) in vmPFC and ERC.

Second, we assessed if 6-fold grid modulation was the best model out of all n-fold grid modulation GLMs. If there is indeed hexagonal grid modulation, a 6-fold model should explain the most variance compared to 4-fold, 5-fold, 7-fold, and 8-fold modulation models. We repeated the grid-modulation GLM analysis in the four ROIs for 4∼8fold regressors: cos(*n*θ_*t*_), sin(*n*θ_*t*_), *n* = 4∼8. Since all the models have the same number of parameters, we compared the 4-fold, 5-fold, 6-fold, 7-fold, and 8-fold models using the z-converted *F*-statistics of the grid angle regressors to assess which set of n-fold regressors explain the most variance in each ROI. Furthermore, to set up clear expectations of the resulting pattern under different hypotheses, we simulated the analysis as close as possible by generating the BOLD signal for each subject and regressing these simulated data with the 4∼8fold regressors. To simulate the results of the analysis when the signal is subjective value, we used the individual’s fitted hyperbolic subjective value as modulators of neural activity that were convolved with a double-gamma HRF with autocorrelated noise added (drawn from a multivariate Gaussian distribution with inter-TR correlation of 0.12). To simulate the results of the analysis when the signal is a hexagonal gridlike modulation, we randomly chose one unique grid angle for each subject (between 0 and 60 degrees) and calculated each trial’s neural activation according to the alignment between the grid angle and trial angle (n=6):

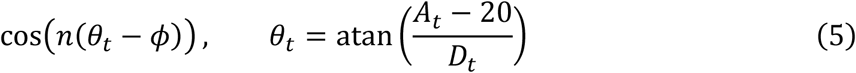

The resulting hexagonal modulation was also convolved with a double-gamma HRF with autocorrelated noise added. The simulated subjective value BOLD signal and the hexagonal grid BOLD signal were then regressed against the same regressors used for the real data.

To ensure the robustness of the results, we performed the model comparison analyses above in alternatively defined ROIs and also in alternatively scaled attribute spaces. For the former, we defined spherical ROIs from peak GLM coordinates closest to the vmPFC and ERC peaks in Constantinescu et al. (2016) and to the vmPFC and VS peaks in Bartra et al. (2013). For the latter, we calculated n-fold grid modulation regressors in an attribute space where both amount and delay were min-max normalized to have equal range. The model comparison results in these alternative ROIs and alternative attribute spaces were the same as in the original ROIs and original attribute spaces and are therefore presented in the supplemental materials.

### fMRI analysis – cross-session consistency analysis

Third, following the methods of previous studies, we tested the consistency of grid angles across session 1 and session 2 data. Based on the properties of grid cell representations in non-human animals, it is assumed that for a given brain region each person’s hexagonal grid is oriented at a unique angle that stays constant across time. For example, when people are navigating through a two dimensional space, one person’s grid may have a 6-fold modulating activity that peaks when the person traverses through space at 20 + 60*x* (*x =* 0 … 5) degree angles while another person’s activity may peak at 40 + 60*x* (*x =* 0 … 5) degree angles. We test for such consistency by first calculating each individual’s unique grid angle from the first session’s data using cos(*n*θ) and sin(*n*θ) as regressors. The average coefficients in each of the four 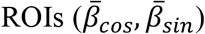 were used to calculate that individual’s *n*-fold grid angle for that ROI 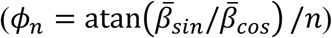. Then, we tested if neural activity in the same ROI in session 2 was aligned with this grid angle by using the consistency regressor shown in eq. 5. The average *z*-statistic of this consistency regressor within the pre-defined ROIs was used to measure the consistency effect across sessions. The key test was whether the consistency effect was the highest at 6-fold, rather than 4-, 5-, 7-, or 8-fold modulation. Again, to set up clear expectations of the results under different hypotheses, we simulated the results of this grid angle consistency analysis under two conditions: when the underlying signal is subjective value, and when the underlying signal is hexagonal modulation. For subjective value, we calculated the 102 participants’ subjective values in both session 1 and session 2 separately via eq. 3 and simulated the BOLD response by convolving the subjective value modulation with a double gamma HRF. For 6-fold modulation, we picked a random angle between 0 and 60 degrees for each subject, which stayed the same across both sessions, and simulated a 6-fold modulation signal according to eq. 5 (with *n* = 6). The resulting simulated neural activity were then convolved with a double-gamma HRF. After adding autocorrelated noise to both subjective value and grid signal, we performed the grid-angle consistency analysis in the same manner as on the real data as outlined above.

The grid angle consistency analysis is the only test that uses data from session 2. As reported in Kable et al. (2017), after session 1 participants were randomized to either a cognitive training intervention or an active control condition. As there were no differences between the two groups in brain activity or decision-making in session 2, we combine them in the analyses reported below. However, when we performed the consistency analysis in the control group only, we found the same results.

## Results

Here we compare two potential coding schemes for vmPFC during decision-making, subjective value and hexagonal grid modulation, both of which are functions of the two attribute dimensions of the choice options. In intertemporal decision-making, the domain we study here, the two attribute dimensions are the amount of the money and the delay until its receipt. A subjective value code predicts that the activity elicited by an option increases as the amount of money increases and decreases as the delay to receipt increases (**Fig. 1a**). The relative slopes of these changes will vary across people depending on the relative weight they place on monetary amounts and delays. On the other hand, a hexagonal gridlike code predicts that activity will vary as a function of the angle in two-dimensional attribute space between the two options that are being compared, with highest activity when this angle matches the person’s unique grid angle (**Fig. 1b**). This prediction assumes that comparing two options in a choice task is akin to conceptually navigating between them in two-dimensional attribute space. Because the hexagonal grid is symmetric, travel in both directions (that is, from option 1 to option 2 or vice versa) yield equivalent predictions. An important constraint in testing these two hypotheses is that only comparisons between two options that involve a tradeoff between the two attributes are non-trivial. That is, most two-attribute choice problems involve a comparison between one option that is better on one attribute and another option that is better on the other attribute; in the intertemporal decision-making case, this translates to choices between greater amounts of money at longer delays versus lesser amounts of money at shorter delays. Both decision theory and neural evidence suggest that dominated choices, in which one option is better than the other on both attributes, can use different psychological and neural mechanisms than choices that involve tradeoffs between the attributes (Hunt et al., 2012; Kahneman & Tversky, 1979). Thus, most two-attribute choice tasks, including the choice task we use here, only sample two quadrants within the two-dimensional attribute space (**Fig. 1a-b**).

In the intertemporal choice task we study, one of the two options is fixed, so that participants choose on every trial between an immediate option that is always $20 and a delayed option that varies from trial to trial in monetary amount and length of delay. This feature of the design simplifies testing for correlates of subjective value, as we can now test for signals modulated by subjective value of the variable, delayed larger reward, according to that individual’s intertemporal preferences (captured by their discount rate for delayed rewards, **Fig. 1c**). However, this feature does not complicate our ability to test for a hexagonal gridlike code; though it is equivalent to restricting the space of choices to one quadrant of Figure 1b, given the symmetry of the hexagonal grid code, it does not further alter or restrict the range of potential grid angles sampled in our design (**Fig. 1d**).

We first show via simulation that a hexagonal grid modulation over decision attribute space could in theory account for previously observed neural correlates of subjective value in this task (Cox & Kable, 2014; Kable et al., 2017; Kable & Glimcher, 2007, 2010). We calculated the best-fitting grid angle for different subjective value landscapes generated assuming different intertemporal preferences (i.e., different discount rates for delayed rewards). **Fig. 2** shows subjective value signals and their best-fitting gridlike signals at various discount rates. The maximum correlation between the two signals ranges between *r* = 0.5 and *r* = 0.7 depending on the discount rate. These high correlations suggest that it is possible for a hexagonal modulation and subjective value to be confused with each other and motivate testing and comparing these two accounts in neural data.

**Figure 2.**
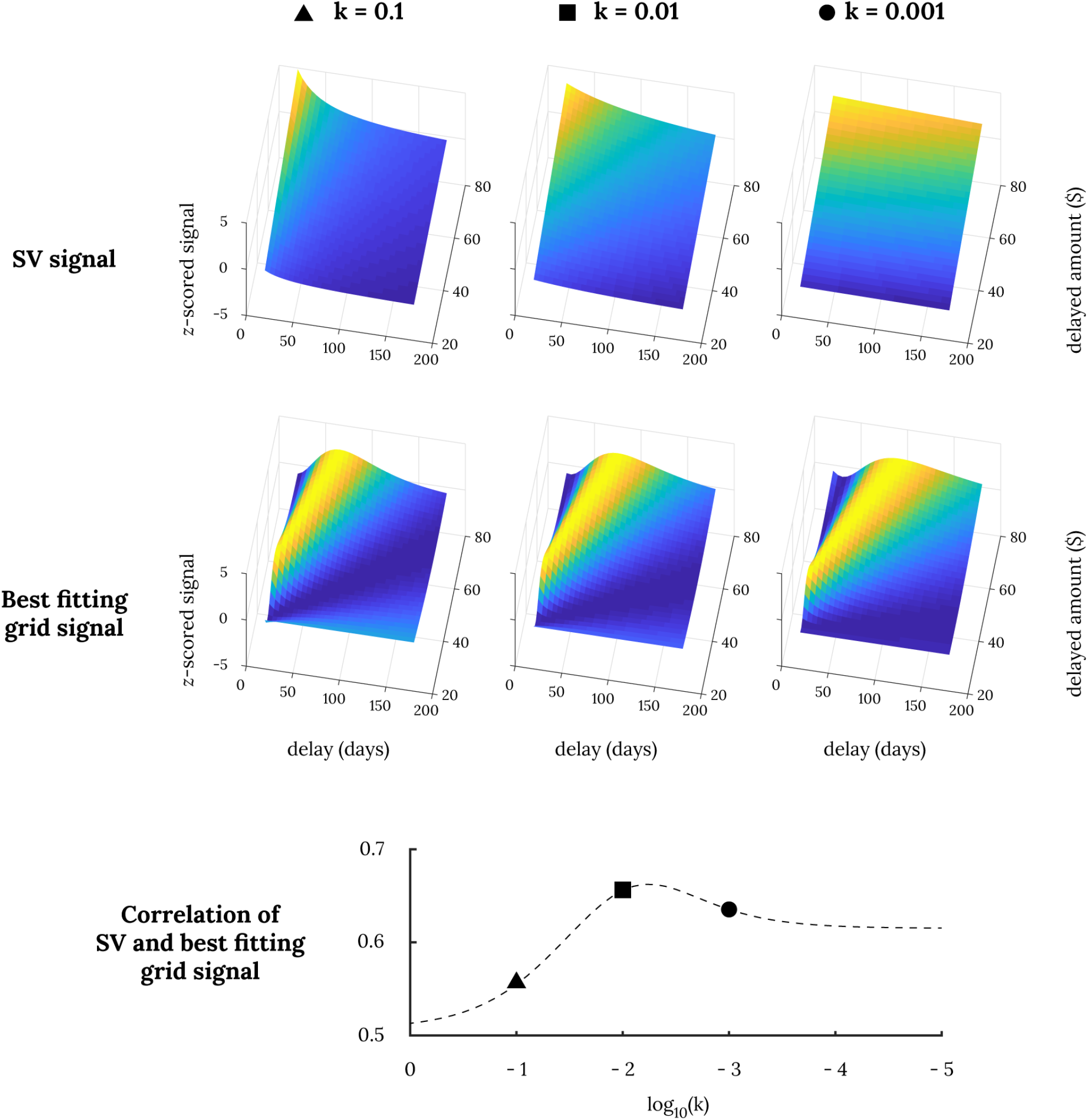
Correlation between SV signal and its most similar hexagonal modulation signal. The top three panels show simulated subjective value signals for various delayed amounts at various discount rates and the next three panels below show their respective best fitting hexagonal grid modulations. The correlations between the two signals are provided below in dotted lines across various discount rates.

We next test for neural activity correlated with subjective value or gridlike modulation in real data. Subjects in our study participated in two imaging sessions separated by ten weeks, and we perform this test in session 1 data. **Fig. 3** shows significant effects across the whole brain for both subjective value and gridlike modulation regressors. Perhaps given the statistical power in our dataset (n = 145), we observed significant effects in widespread brain regions for both analyses. The subjective value effects, which we have reported previously for this dataset (Kable et al., 2017), include peaks in the vmPFC and ventral striatum that overlap with previous meta-analyses of subjective value correlates (Bartra et al., 2013). The gridlike modulation effects include peaks in the vmPFC and entorhinal cortex (ERC) as observed previously during conceptual navigation by Constantinescu et al. (2016).

**Figure 3.**
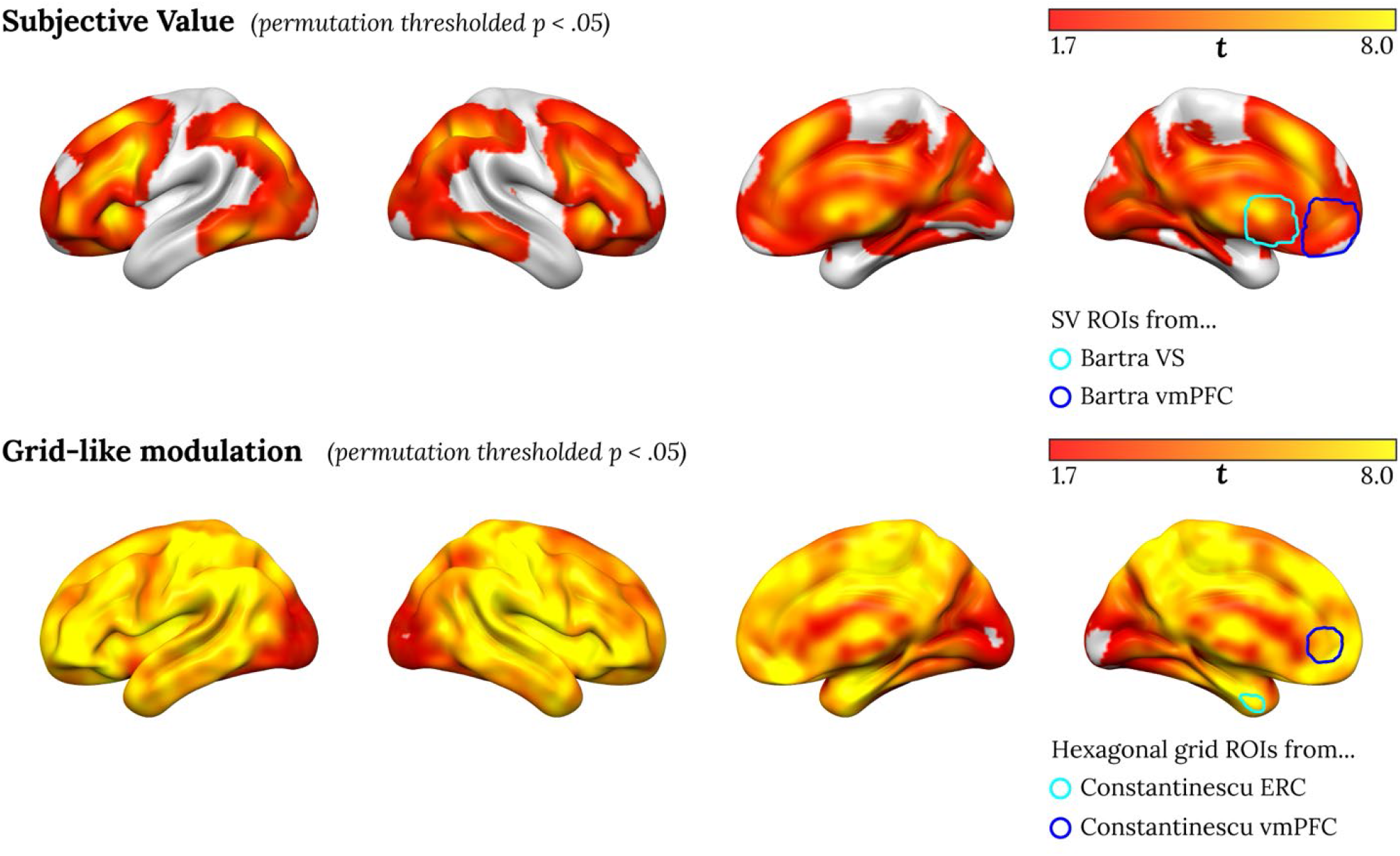
Regions with significant subjective value correlation (top) and significant hexagonal gridlike modulation (bottom) in session 1. (*p* < 0.05 with permutation testing with threshold free cluster enhancement). The rightmost brain shows overlays of the two ROIs from Bartra et al. (2013) on top and the two ROIs from Constantinescu et al. (2016) on bottom.

The widespread effects observed for the gridlike modulation analysis, which encompass almost the entire brain, emphasize an important point: given that the angle of grid modulation is a free parameter, even between runs, these regressors provides a good degree of flexibility to fit a diversity of response patterns. Hence, definitive evidence for a hexagonal grid modulation signal requires showing that modulation is stronger at 6-fold than at other folds and that the angle of grid modulation is consistent across time. We turn to these stronger tests as we next employed three different ways to directly compare subjective value and hexagonal gridlike codes as accounts of neural activity in this task.

First, we simply compare the AICs of the subjective value and hexagonal grid GLMs above. We found the subjective value GLM was a better descriptive model of fMRI activity in all four ROIs tested (**Fig. 4**). Most subjects had smaller AICs for the subjective GLM than for the hexagonal grid GLM in both the vmPFC and ventral striatum ROIs drawn from Bartra et al. (2013), as well as in both the vmPFC and ERC ROIs drawn from Constantinescu et al. (2016). We also performed this comparison across all voxels in the brain, and we did not find any voxels where the hexagonal grid GLM has significantly lower AIC.

**Figure 4.**
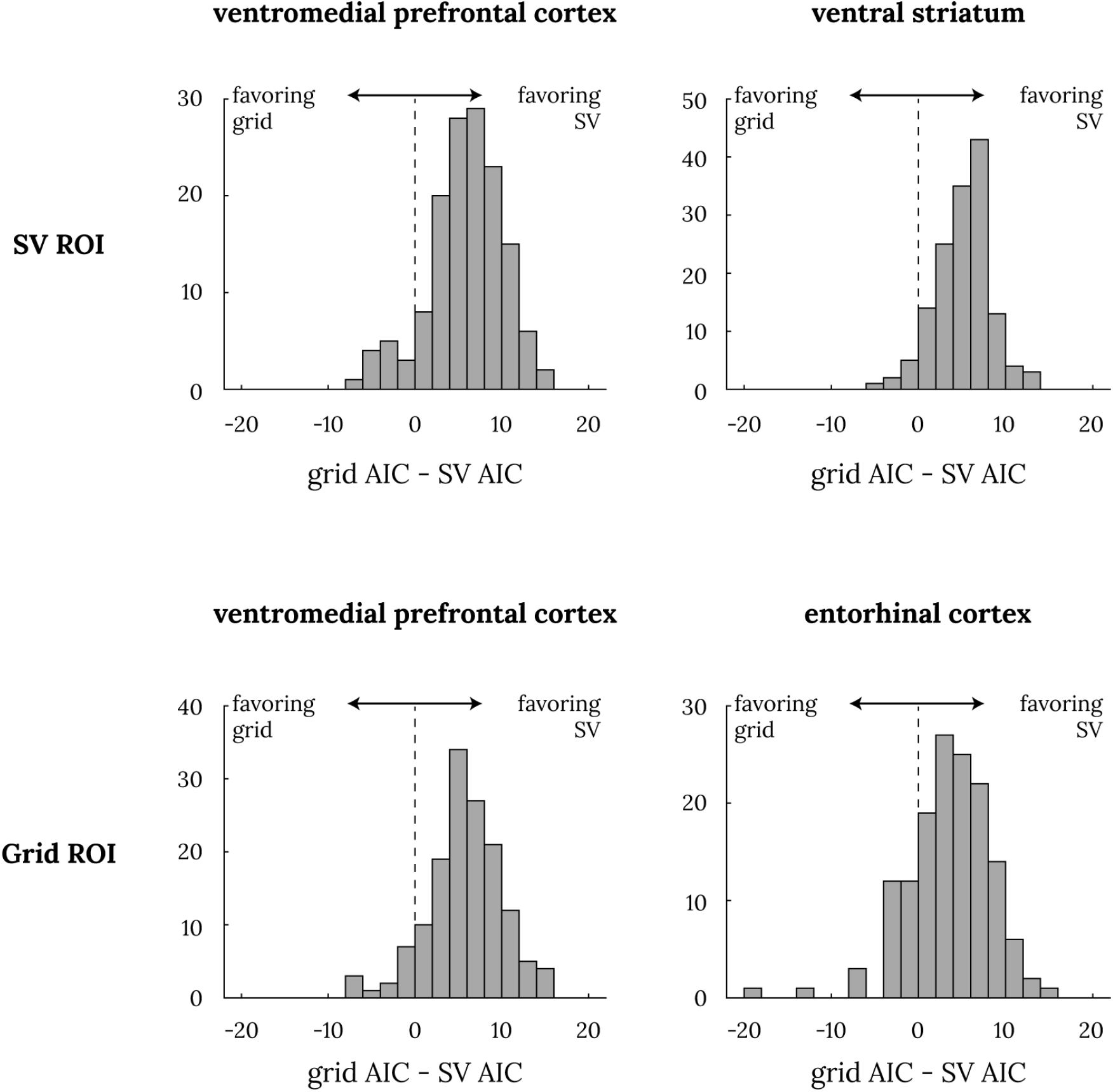
Model comparison between subjective value GLM and hexagonal grid GLM. Each of the four panels show the histogram of individual AIC differences between the hexagonal grid GLM and the subjective value GLM such that a positive number indicates AIC difference favoring the SV model. The top two panels are from the ROIs from Bartra et al. (2013); the bottom two panels are from the hexagonal grid ROIs from Constantinescu et al. (2016). All four ROIs’ mean AIC difference was significantly different from 0 at *p* ≪ .001 with a one-sample t-test.

Despite this significant result in all four ROIs, one may wonder if the subjective value GLM is favored simply because AIC favors models with fewer parameters. Our next test avoids this issue, as we compare different n-fold grid modulation GLMs in the same ROIs. If the true signal is a hexagonal gridlike modulation, a 6-fold modulation model should account for more variance in the data compared to 4, 5, 7, or 8-fold modulation (**Fig. 5**). In contrast, if the true signal is subjective value, 4-fold modulation should account for the most variance, and variance explained should decline across 4∼8 folds (**Fig. 5**; in other words, 4-fold modulation best mimics a subjective value signal, which in our task is always highest in one corner of the two-dimensional attribute space and lowest in the opposite corner). In none of the four ROIs did we find that 6-fold modulation was not the best descriptor of the BOLD signal. In fact, in both the vmPFC and ventral striatum ROIs from Bartra et al. (2013), and in the vmPFC ROI from Constantinescu et al. (2016), we found that 4-fold modulation explains significantly more variance than the 6-fold modulation (**Fig. 5**). This difference was not significant in the ERC ROI from Constantinescu et al. (2016). Again, we also performed this comparison across all voxels in the brain, and we did not find any where the 6-fold model explains significantly more variance than the 4-fold model.

**Figure 5.**
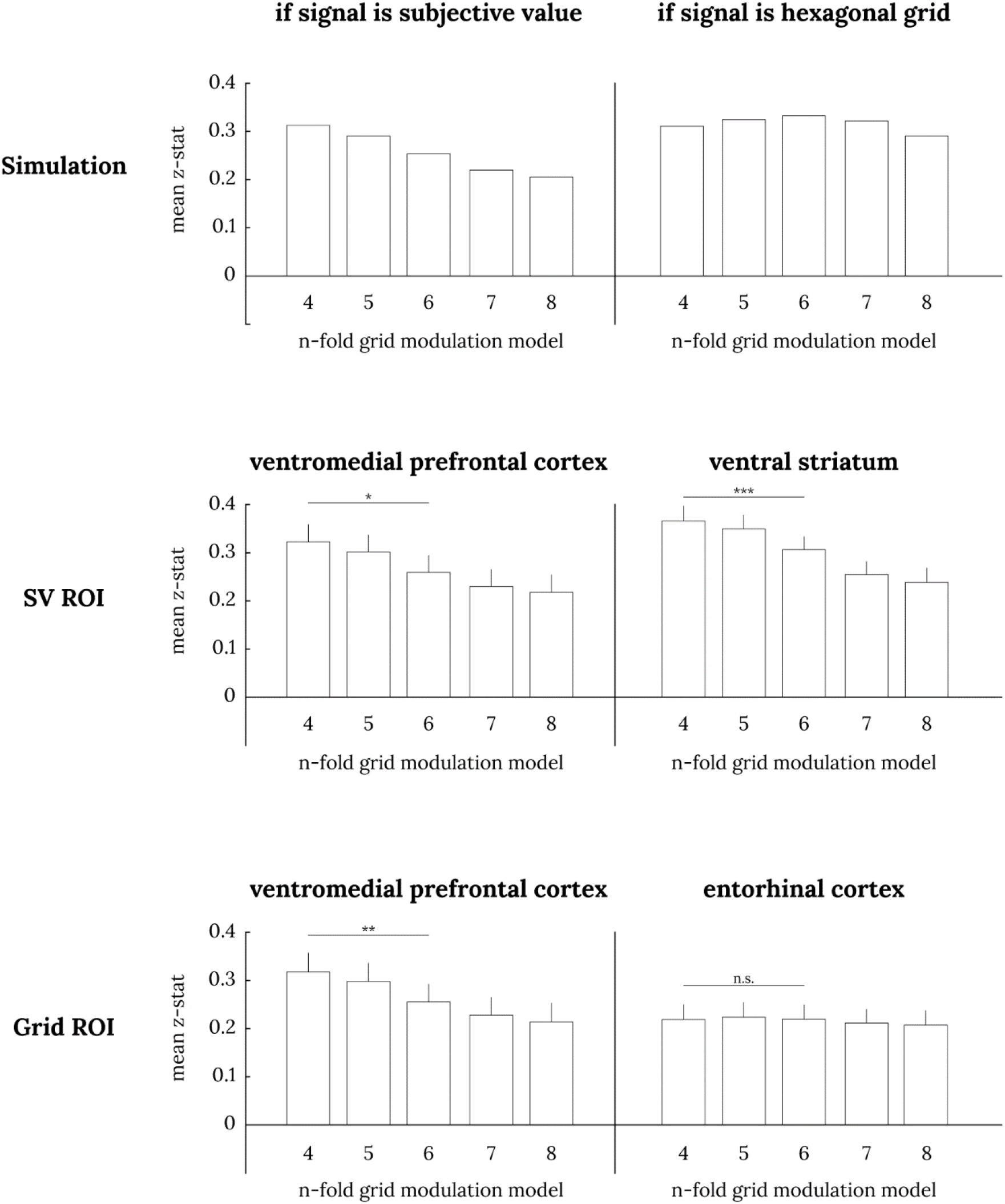
Comparison of n-fold grid modulation GLMs in session 1 data. The top two panels show simulated results of n-fold grid modulation GLMs when the true signal is SV (left) or a hexagonal gridlike code (right). When the true signal is subjective value, we expect a descending staircase pattern, and when the true signal is a hexagonal grid, we expect a pyramid pattern. The middle two panels show the GLM analysis in SV ROIs from Bartra et al. (2013); the bottom two panels show them for hexagonal grid ROIs from Constantinescu et al. (2016). The error bars denote the standard errors of the mean. Paired t-test between 4-fold and 6-fold: * *p* < .05, ** *p* < .01, *** *p* < .001.

As a third test, we take advantage of the fact that subjects participated in two sessions in our study, and perform the exact grid angle consistency analysis proposed to be the critical test for gridlike responses in Constantinescu et al. (2016). This test fixes the grid angle based on the fit of 6-fold modulation GLM in the first session and tests for modulation at that grid angle for 4∼8 fold-modulation in the second session. Similar to the analysis above, if the true signal is hexagonal modulation, we should see the strongest grid-angle consistency effect with 6-fold modulation. In contrast, if the true signal is subjective value, we should see the strongest grid-angle consistency effect with 4-fold modulation. This analysis of grid angle consistency analysis again unambiguously shows that vmPFC activity is consistent with subjective value and does not exhibit consistent hexagonal gridlike modulation (**Fig. 6**). We found that the grid-angle consistency effect was significantly larger at 4-fold modulation than 6-fold modulation in both the vmPFC ROIs from Bartra et al. (2013) and from Constantinescu et al. (2016).

**Figure 6.**
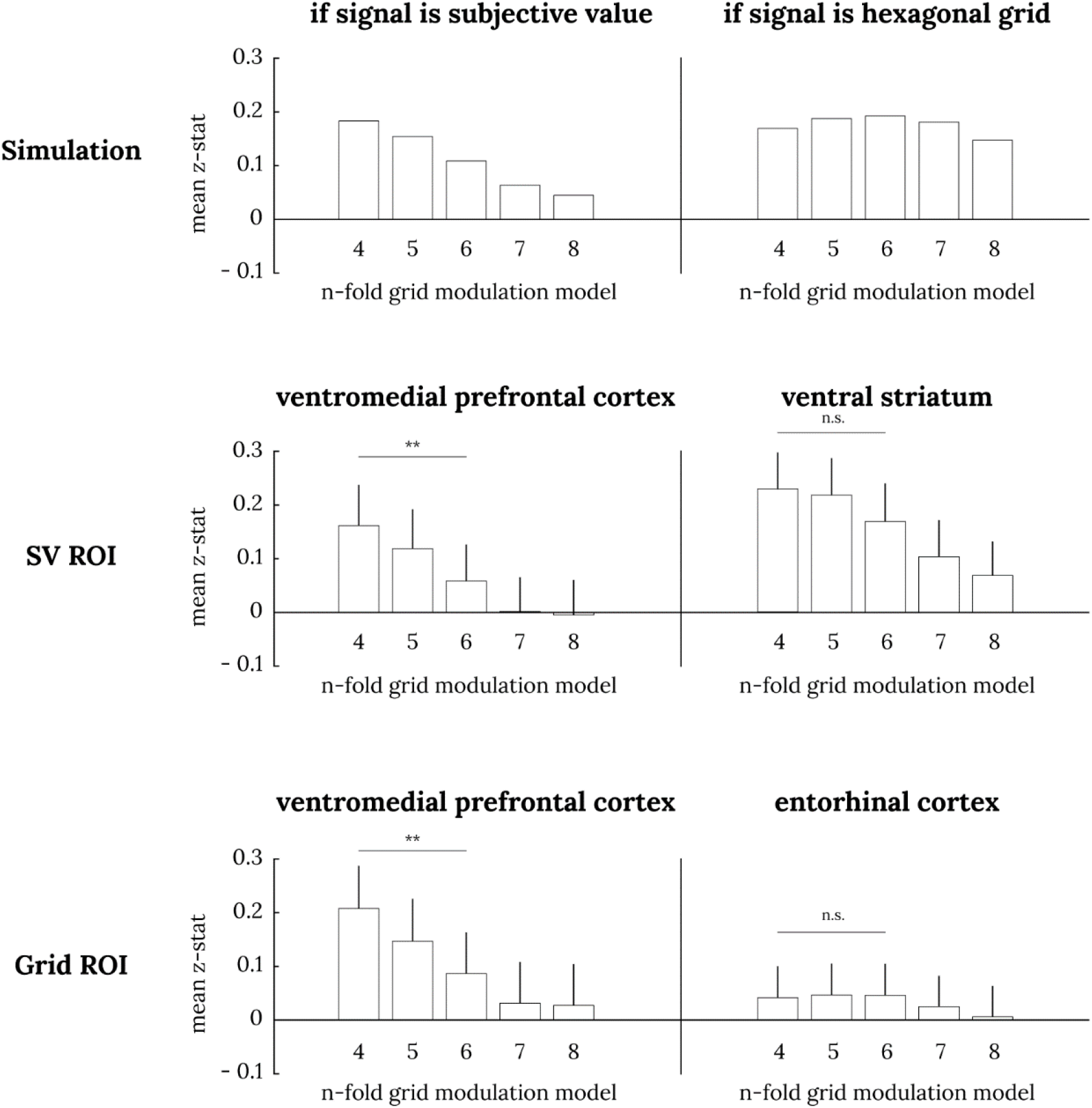
Grid-angle consistency analysis in session 2 data. The top two panels show simulated results of grid-angle consistency analysis when the true signal is subjective value (left) or a hexagonal gridlike code (right). When the true signal is subjective value, we expect a descending staircase pattern, and when the true signal is a hexagonal grid, we expect a pyramid pattern. The middle two panels show the grid-angle consistency analysis in subjective value ROIs from Bartra et al. (2013); the bottom two panels show them for hexagonal grid ROIs from Constantinescu et al. (2016). Paired t-test between 4-fold and 6-fold: ** *p* < .01.

Finally, we perform several additional analyses to demonstrate the robustness of these results under different sets of assumptions. First, our ROIs, defined based on past studies, could have missed the exact location of gridlike modulation in the current study. If we define subjective value and gridlike code ROIs based on the peaks of these analyses in the current study, rather than on previous studies, we observe exactly the same pattern of results (**Supplementary Fig. 1**). Second, our gridlike modulation analyses assume that in the scaling of two attribute dimensions one dollar is equivalent to one day. If we alternatively assume that the two dimensions are such that the range of amounts and delays sampled cover an equivalent scale, we again observe exactly the same pattern of results as above (**Supplementary Fig. 2**).

## Discussion

During decision-making, vmPFC activity has previously been shown to correlate with subjective value; here we tested whether it could instead be better explained by hexagonal gridlike activity reflecting conceptual navigation through the space defined by the attributes of the choice options. Theoretically, this idea is plausible as we found that there can be a high correlation between putative subjective value and hexagonal gridlike modulation signals. Empirically, we tested for gridlike responses in vmPFC in a large fMRI dataset of a standard two-attribute intertemporal decision task involving two sessions. We found, unambiguously across three different tests, that the vmPFC signal during intertemporal decision making is better explained by a subjective value than a hexagonal gridlike signal. First, a simple model comparison between a GLM that assumed activity was correlated with subjective value and one that assumed activity was modulated in a hexagonal gridlike manner favored the subjective model in vmPFC. Second, across various n-fold grid modulation models, the 6-fold hexagonal modulation model was not the best model of vmPFC activity. Rather, a 4-fold modulation that could best mimic subjective value was the best fitting model. Thirdly, we found that the cross-session consistency of an individual’s grid angles was not the highest when assuming 6-fold hexagonal modulation. Instead, we again found that the 4-fold modulation model resulted in higher cross-session grid angle consistency, matching our simulation results for a subjective value signal. Thus, we found strong evidence that BOLD activity in vmPFC during intertemporal decision-making is correlated with subjective value and does not reflect a hexagonal gridlike code.

Though there are assumptions in any one of the analyses we performed, the strong convergence of results across analyses and robustness checks lends strength to these conclusions. For example, the direct model comparison requires a choice about how to penalize the hexagonal grid model for its greater number of parameters, the grid model requires an assumption about the relative scaling of the two attribute dimensions, and the grid angle consistency analysis assumes stability of this representation across ten weeks. (In the last case, ten weeks is longer than has previously been demonstrated in humans or other animals, though subjective value correlates are consistent across ten weeks (Kable et al., 2017), so for a gridlike code to explain these it would require grid angle consistency over this delay.) However, across three different kinds of tests, for two different methods of defining ROIs and for two different assumptions about the scaling of attribute dimensions, we consistently find evidence consistent with a subjective value signal and no significant evidence at all for a hexagonal gridlike code in vmPFC.

Our results constrain the implications of Constantinescu et al. (2016) by limiting the conditions under which a gridlike code is observed in the vmPFC. Our intertemporal choice task is obviously very different from the sort of mental navigation demanded in Constantinescu et al. (2016). Their study involved extensive learning of a novel conceptual space by the participants prior to the navigational task; ours takes advantage of a spontaneous conceptual space created by the option attributes as they are presented in a choice task. Their navigational task involved a period of mental simulation where the participants are asked to imagine the progress of the stimulus, akin to virtual navigation, whereas we simply ask participants to make a choice. Nevertheless, the two tasks share important similarities in that both operate within a two-dimensional space where gridlike representations of the task structure might be expected and indeed have been proposed (Bellmund et al., 2018). Our findings suggest that the two-dimensional space defined by the option attributes that is available during decision-making does not necessarily provoke gridlike representations.

The vmPFC is important for a wide variety of functions, from learning and decision-making to schematic memory and social cognition (Fehr & Camerer, 2007; Gilboa & Marlatte, 2017; Grabenhorst & Rolls, 2011; Janowski, Camerer, & Rangel, 2013; Lieberman, Straccia, Meyer, Du, & Tan, 2019; Powell & Redish, 2016; Roy, Shohamy, & Wager, 2012). A possible explanation for this diversity is that different subregions of the vmPFC serve different functions. Hence, a strong possibility is that the vmPFC represents the *relevant* cognitive map for the current task (Bernardi et al., 2018; Schuck, Cai, Wilson, & Niv, 2016; Wilson, Takahashi, Schoenbaum, & Niv, 2014), and therefore the nature of the coding scheme in vmPFC may depend on the demands of the task at hand. A decision-maker does not necessarily need to know the spatial relationships between different choice options in attribute space but does need to know the relevant ordering of different options in terms of her preferences. Subjective value is a representation that has long been known to afford this useful feature for decision-making; using such a representation of multi-attribute options and choosing the maximal valued option is guaranteed to result in transitive, non-cyclical choices (Samuelson, 1937). Subjective value may therefore be the more relevant representation of the option space for the kind of decision tasks we studied here.

## Acknowledgements

This study was funded by grants from the National Cancer Institute R01 CA170297 (JK, CL) and R35 CA197461 (CL)

## Author Contributions

Conceptualization, L.Q.Y; Methodology and Formal Analysis, S.L; Investigation, all authors; Writing - Original Draft, S.L. and L.Q.Y.; Writing Review & Editing, all authors; Supervision J.W.K.; Funding Acquisition C.L., and J.W.K.

## Declaration of Interests

The authors declare no competing interests.

## Notes

### Competing Interest Statement

The authors have declared no competing interest.

