## supplemental materials for "Subjective value, not a gridlike code, describes neural activity in ventromedial prefrontal cortex during value-based decision-making"

*Model comparisons using ROIs generated from our dataset*

To address the potential concern that ROIs from previous studies (i.e., Bartra et al. (2013) ROIs for subjective value and Constantinescu et al. (2016) ROIs for hexagonal grid) may not exactly match the loci of activation in our data, we defined additional ROIs based on session 1 GLM peak statistics for subjective value and hexagonal grid regressors. For the subjective value GLM, we created a spherical ROI around the peak for the permutation $t$-score for the subjective value regressor that was closest to the center of the Bartra et al. (2013) ROIs. For the hexagonal grid GLM, we created a spherical ROI around the peak $z$-transformed $F$-statistic for the grid angle regressors that was closest to the center of the Constantinescu et al. (2016) ROIs. We found that the SV ROIs based on $t$-statistics were entirely encapsulated by the ROIs from Bartra et al. (2013), while the hexagonal grid ROIs based on $F$-statistics were shifted slightly from the ROIs from Constantinescu et al. (2016) (**Fig S.1a**).

Model comparison results in these ROIs were identical to those reported in the main manuscript. In all four ROIs, the subjective value GLM had significantly lower AIC scores at the group level (one-sample t-test $p \ll .001$ for all four ROIs; **Fig S.1b**). Furthermore, in all four ROIs, the 6-fold modulation GLM was not the best out of all n-fold modulation models (**Fig S.1c**). In both vmPFC ROIs and the ventral striatum ROI, the 4-fold modulation GLM explained significantly more variance than the 6-fold GLM at the group level, as expected if the true signal is subjective value.

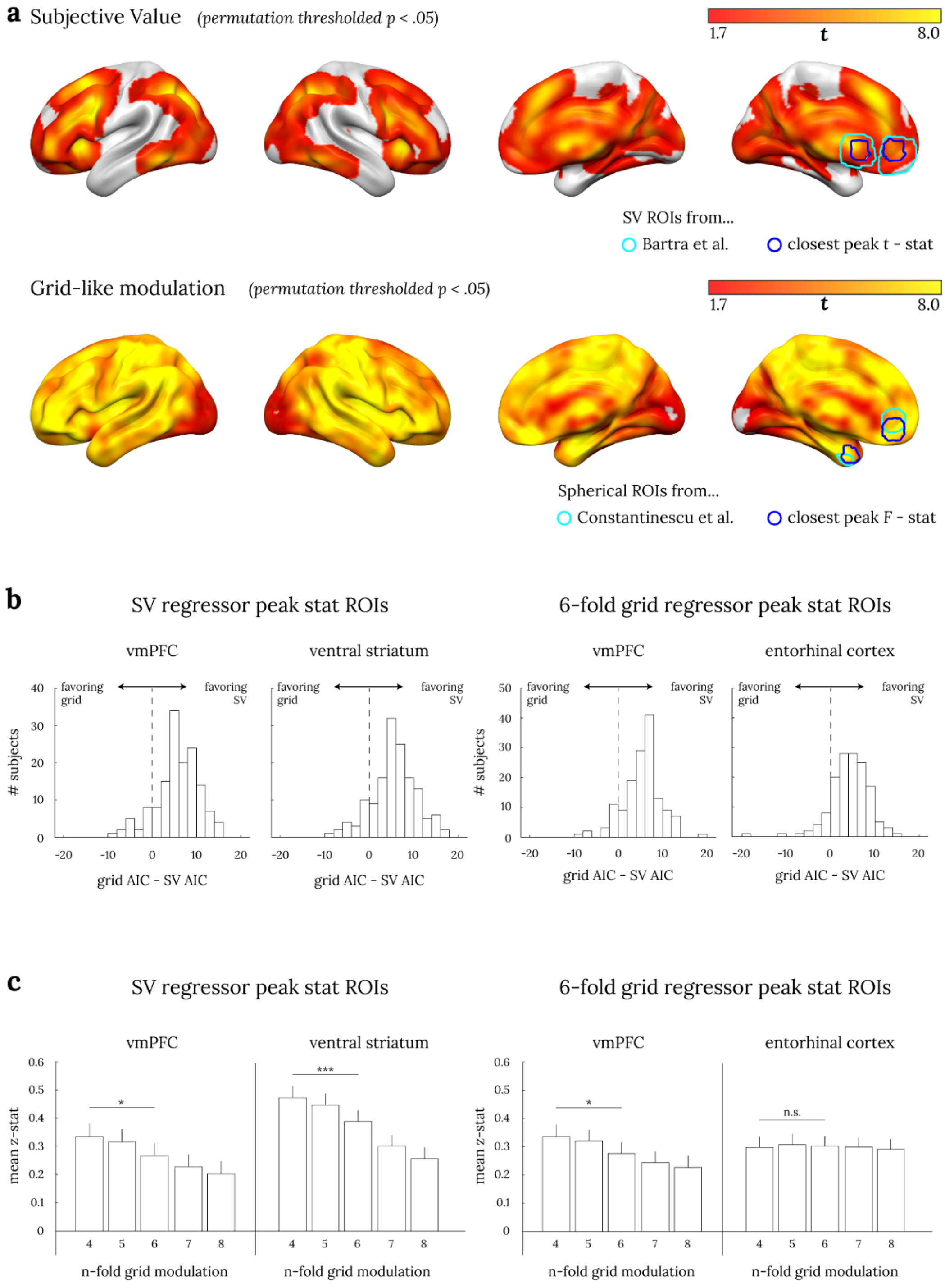


**Fig. S.1. GLM-based ROIs (a) and their model comparison results (b, c).** Panel a shows ROIs defined from peak coordinates of subjective value and hexagonal signal GLMs, compared with ROIs from previous research. Panel b shows the AIC model comparison between the subjective value GLM and hexagonal grid GLM in the four ROIs defined in panel a. Panel c shows the model comparison between various n-fold grid GLMs in the four ROIs defined in panel a. The error bars are standard errors of the mean. Paired t-test between 4 fold and 6 fold: * $p < .05$, *** $p < .001$.

*Alternatively scaled attribute space*

To address whether our results depend on particular assumptions about the scaling of each attribute (i.e., amount and delay) dimension, we re-performed the model comparison analyses using n-fold modulation GLMs in a min-max normalized attribute space, where both the amount and the delay have been normalized to have equal maximum distance. Compared to the original attribute space where delay had a wider range (0~180 days) relative to amount (20~85 dollars), this normalized space treats both variables equally, thereby slightly affecting the traversing angle calculations.

Even in this alternatively scaled space, however, the comparison results were substantively the same as those from previous analyses: the subjective value model was a better descriptor of BOLD activity than the hexagonal modulation model in the four ROIs (**Fig S.2a**, one-sample t-test $p \ll .001$ for all four ROIs), and 4-fold modulation models were better descriptors of BOLD activity than 6-fold modulation models in both vmPFC ROIs and the ventral striatum ROI (**Fig S.2b**). Of note is that the expected pattern of 4~8fold modulation is slightly different from the previously expected pattern when the subjective value is the true underlying signal: from simulation, we expect that 7-fold is the worst model and that 8-fold model is roughly similar 6-fold. This exact pattern of data was found in BOLD activity as well, which provides strong support that the true signal is subjective value.

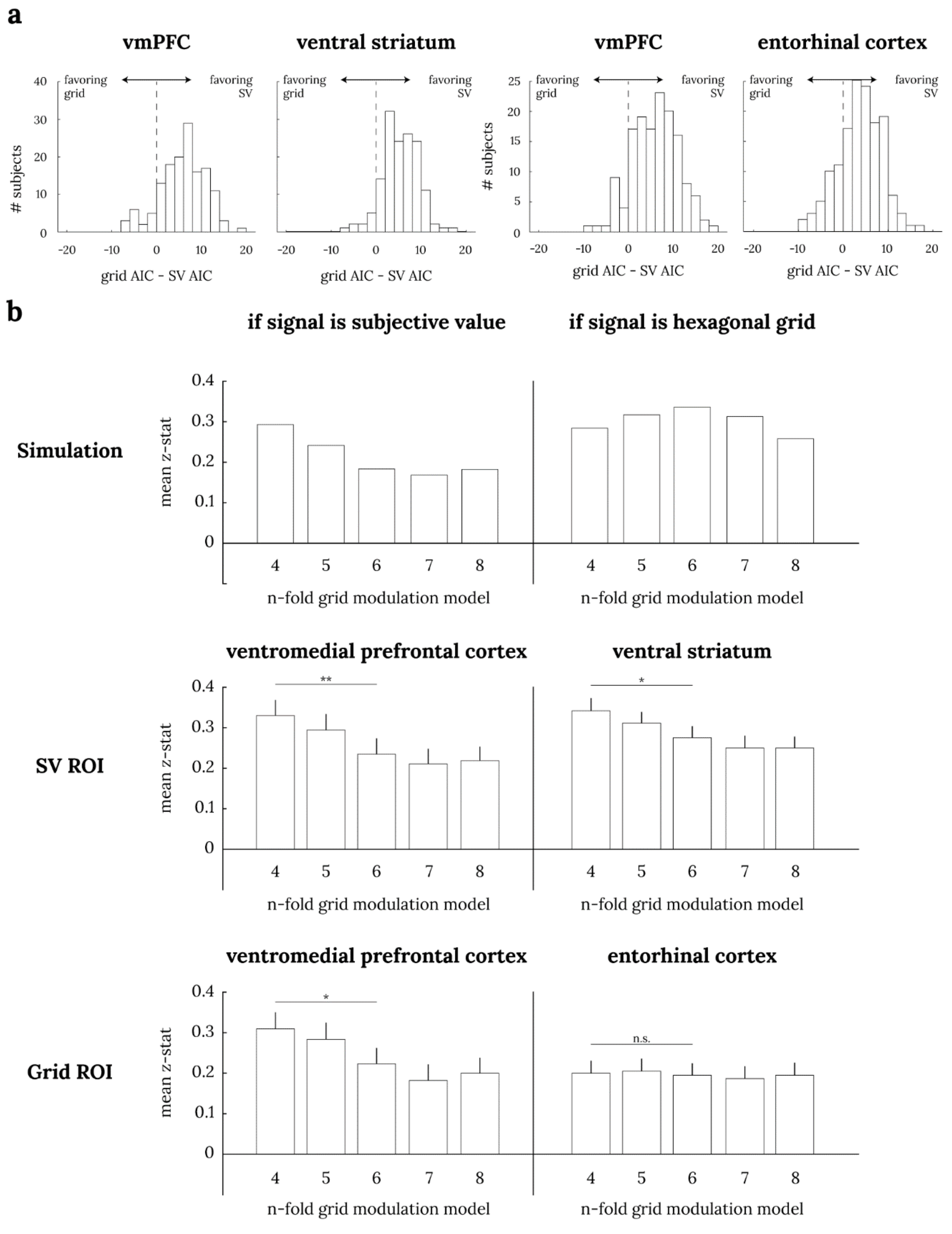


**Fig. S.2. Model comparison results based on min-max normalized attribute space grid modulation GLM.** Panel a shows the AIC model comparison between the subjective value GLM and the hexagonal grid GLM in a new attribute space where delay and amount are min-max normalized to be equal in distance. Panel b shows the model comparison between various n-fold grid GLMs in this min-max normalized space. The error bars are standard errors of the mean. Paired t-test between 4 fold and 6 fold: * $p < .05$, ** $p < .01$.
